# Negative feedback equalizes polarity sites in a multi-budding yeast

**DOI:** 10.1101/2024.12.28.630626

**Authors:** Alex W. Crocker, Claudia A. Petrucco, Kaiyun Guan, Alison C.E. Wirshing, Joanne L. Ekena, Daniel J. Lew, Timothy C. Elston, Amy S. Gladfelter

## Abstract

Morphogenesis in fungi and animals is directed by polarization of small GTPases Cdc42 and Rac. In the budding yeast *Saccharomyces cerevisiae* competition between polarity patches results in one polarized patch and the growth of a single bud. Here, we describe cell polarity in the yeast *Aureobasidium pullulans*, which establishes multiple coexisting polarity sites yielding multiple buds during a single cell division cycle. Polarity machinery components oscillate in their abundance in these coexisting sites but do so independently of one another, pointing to a lack of global coupling between sites. Previous theoretical work has demonstrated that negative feedback in a polarity circuit could promote coexistence of multiple polarity sites, and time-delayed negative feedback is known to cause oscillations. We show that both these features of negative feedback depend on a protein we identified as Pak1, and that Pak1 requires Rac1 but not Cdc42 for its localization. This work shows how conserved signaling networks can be modulated for distinct morphogenic programs even within the constraints of fungal budding.

## INTRODUCTION

The diverse shapes and motile behaviors of eukaryotic cells emerge through the localized assembly and contractility of cytoskeletal elements. Cytoskeletal behavior is orchestrated by a conserved family of Rho GTPases, which become active at specific regions of the plasma membrane^1^. A subset of these, including Cdc42, Rac, and Rop GTPases, are master regulators of cell polarity and serve to orient cell protrusions^2,3^. For many cells, including migratory animal cells, most budding yeasts, and plant pollen tubes, it is critical to develop a single polarity axis^4–6^. However, other cells including neurons^7^ some hyphal fungi^8,9^, and plant xylem precursor cells^10^, develop multiple polarity sites resulting in more complex cell morphologies. How each cell’s polarity machinery is adjusted so as to produce an appropriate number of polarity sites is poorly understood.

The regulatory circuits that control polarity establishment are best understood in the model yeasts *Saccharomyces cerevisiae* and *Schizosaccharomyces pombe*^11^. In these cells a single polarity GTPase, Cdc42, switches between dispersed (unpolarized) and clustered (polarized) states. Clustering is driven by a core positive feedback loop in which active GTP-bound Cdc42 at the plasma membrane recruits a complex containing a Cdc42-directed guanine nucleotide exchange factor (GEF) from the cytoplasm^12–15^. The GEF activates neighboring Cdc42, leading to a local accumulation of GTP-Cdc42 at the plasma membrane. Whereas active GTP-Cdc42 diffuses slowly at the membrane, most inactive GDP-Cdc42 is present in the cytoplasm in complexes with a guanine nucleotide dissociation inhibitor (GDI), and these complexes diffuse much more rapidly^16^. Over time, accumulation of active Cdc42 and GEF complexes at the polarity site leads to partial depletion of inactive Cdc42 and GEF complexes from the cytoplasm, slowing the growth of the cluster. In addition, GTPase activating proteins (GAPs) promote GTP hydrolysis by Cdc42 at the membrane, leading to GDI binding and dissociation of inactive Cdc42 back to the cytoplasm. As more GTP-Cdc42 accumulates in the cluster, Cdc42 inactivation accelerates while further Cdc42 activation slows due to depletion of cytoplasmic species, leading to a balanced steady state^17^. This network of interactions is considered the core positive feedback loop that promotes the establishment of polarity sites.

Computational models of the core positive-feedback circuit described above revealed that when parameters were adjusted to reflect the conditions of *S. cerevisiae*, simulations always led to a single polarity site at steady state^14,18,19^. This was so even when molecular-level fluctuations were included in the model and seeded the initial formation of multiple polarity sites^20,21^. Model analyses showed that as clusters grow large enough to partially deplete cytoplasmic species, they engage in a competition such that Cdc42 and GEF complexes released from small clusters tend to be recruited by larger clusters, until only one large cluster exists. Rapid imaging of polarity establishment in *S. cerevisiae* cells confirmed that cells often developed transient multi-cluster intermediates that appeared to compete with each other, yielding a single winner that went on to bud^22^. Experimental manipulations predicted to slow the competition process led to formation of multi-budded cells^6^, suggesting that a minimal Cdc42 positive feedback circuit can support development of one and only one polarity site in a specific biophysical regime.

More recent theoretical studies have suggested several ways in which the polarity circuit could be modified so as to produce more than one polarity site. First, simply increasing polarity factor concentrations can slow competition and render coexisting polarity sites more stable^19^. Maintaining similar concentrations but increasing the total amounts of polarity factors by making cells sufficiently large can have the same effect^19^. Second, relaxing the mass-conservation assumption and allowing rapid synthesis and degradation of polarity proteins could insulate clusters from competition, potentially leading to multipolar outcomes^23^. Third, addition of a global negative feedback loop to the core circuit could promote equalization, rather than competition, between polarity sitess^22,24–26^. Fourth, addition of localized factors (such as “Bud site selection” proteins in *S. cerevisiae* or “Tip factors” in *S. pombe*) that promote local Cdc42 activation can stabilize clusters against competition at those locations^3,27,28^. Interestingly, although the *S. cerevisiae* polarity circuit does respond to localized factors^29^ and contains negative feedback^22,30,31^, neither of those features leads to multipolarity. However, manipulations that result in cell enlargement and increased expression of polarity proteins can lead to multipolar outcomes due to stalled competition^25^. Whether any of these mechanisms explains multipolar behaviors in cells that naturally make multiple polarity sites is currently unknown.

Here, we investigated cell polarity in *Aureobasidium pullulans*, a morphologically plastic fungus isolated from environments as varied as plant leaves, solar salterns, and arctic glaciers^32,33^. In its budding yeast form, *A. pullulans* cells can be multinucleate and grow a variable number of buds in a single cell division cycle^34^ (Figure 1A). Recent advances have made *A. pullulans* a tractable model for cell biological investigation^35,36^, providing an opportunity to study how its polarity circuit might differ from that of *S. cerevisiae* to produce multipolar outcomes. Our findings suggest that multipolarity is correlated with increased cell size, and that equalization of polarity sites depends on a negative feedback loop involving a GTPase effector kinase, Pak1.

**Figure 1.**
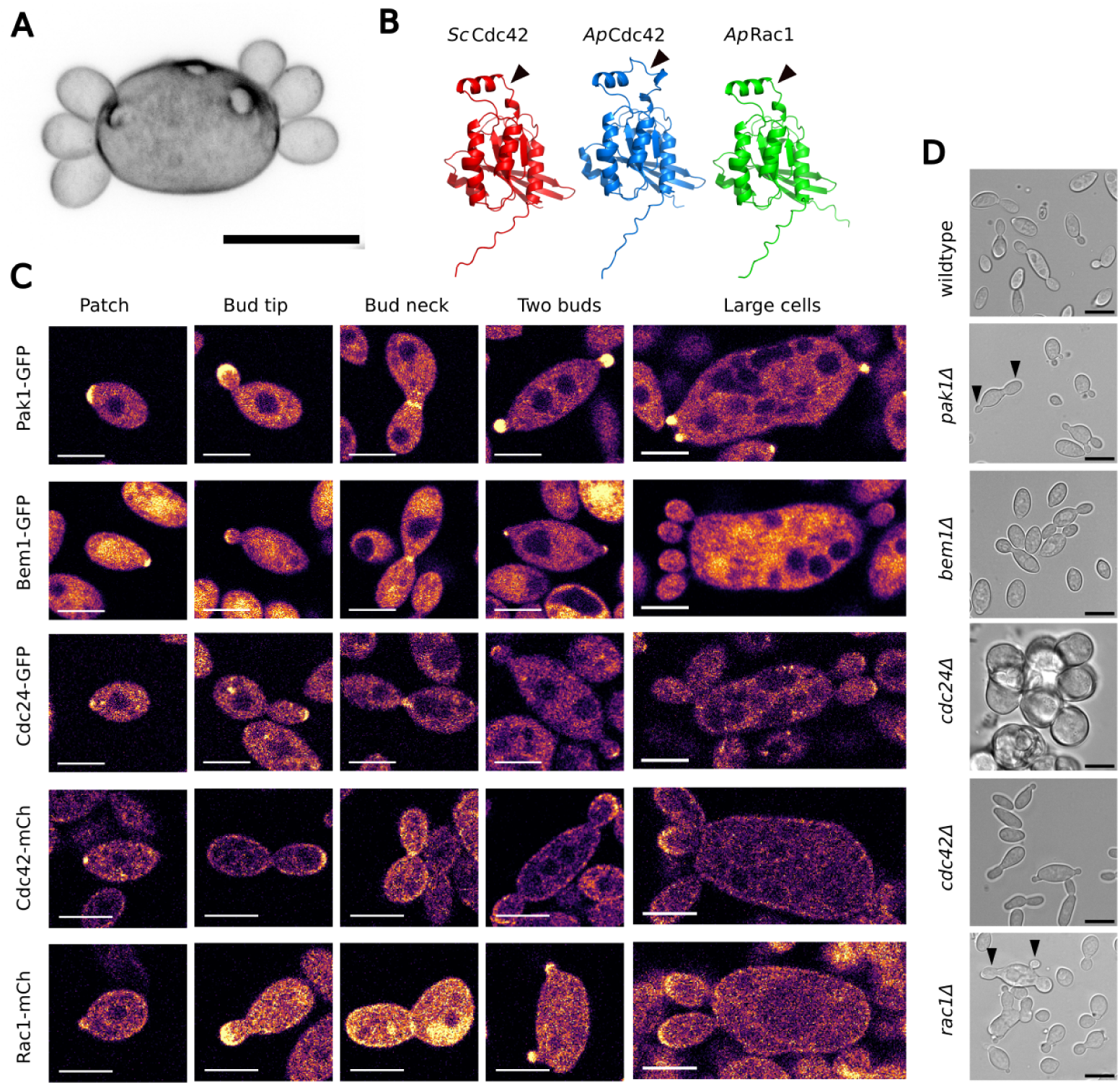
Cell polarity proteins in *A. pullulans*. (A) Image of calcofluor stained *A. pullulans* cell with multiple buds. (B) Predicted structures of Cdc42 and Rac1 proteins. Arrow indicates the location of mCherry tags. (C) Tagged proteins localize to patches, bud tips, and bud necks. Localization of probes is similar in small and large cells. Scale bar, 5 μm. (D) Brightfield microscope images of wildtype and mutant strains deleted for individual polarity proteins.

## RESULTS

### Core polarity proteins are conserved in *A. pullulans* and localize to bud sites

The core polarity circuit in *S. cerevisiae* consists of a single GTPase, Cdc42, which is regulated by a GEF (Cdc24), a GDI (Rdi1), and at least four GAPs (Bem2, Bem3, Rga1, and Rga2). Positive feedback is mediated by two effector p21-activated kinases (PAKs) together with a scaffold protein (Bem1) that binds to the GEF, GTP-Cdc42, and the PAKs^12,13,37–39^. We identified homologs of all of these polarity proteins in *A. pullulans* (Table S1, Figure S1). Compared to *S. cerevisiae*, *A. pullulans* has more GTPases (two instead of one) but fewer GAPs (two instead of four) and PAKs (one instead of two). Based on sequence comparisons, we called the *A. pullulans* genes *CDC42* and *RAC1* (GTPases), *RDI1* (GDI), *CDC24* (GEF), *BEM3* and *RGA1* (GAPs), *PAK1* (PAK), and *BEM1* (Scaffold).

To assess the intracellular localization of the *A. pullulans* polarity proteins, we tagged proteins expected to localize to polarity sites with genetically-encoded fluorophores. Endogenous Pak1, Bem1, and Cdc24 were tagged at their C-termini with a version of GFP that was codon optimized for *A. pullulans*^36^. Since tagging of Cdc42 at either terminus can interfere with its function and localization, we tagged Cdc42 and Rac1 at an internal loop that allows a less perturbing tag in other systems^15,40^ (Figure 1B). The mCherry-tagged versions of Cdc42 and Rac1 were each incorporated, along flanking sequences, as additional copies of the genes at the *URA3* locus^36^. Each of the five tagged proteins was concentrated at distinct foci, including the tips of smaller buds and the mother-bud-necks of larger buds (Figure 1C).

We next investigated the effects of disrupting these genes. Each gene was deleted by replacing the full coding sequence with a hygromycin-or nourseothricin-resistance cassette^35^. All deletion mutants were viable (Figure S2), but deletion of *CDC24* resulted in a severe growth defect. *cdc24*Δ cells did not bud, but rather grew isotropically to a large size before dividing by fission (Figure 1D). Deletion of either *CDC42* or *RAC1* resulted in milder growth defects (Figure S2), with both mutants continuing to bud. Deletion of *PAK1* resulted in a similar phenotype, whereas deletion of *BEM1* did not result in an obvious growth defect. Furthermore, both the *rac1*Δ and the *pak1*Δ mutant mother cells often made buds with unequal sizes (Figure 1D), a phenotype explored in more detail below. In summary, only the GEF was essential for budding, while the other polarity proteins may contribute to budding in a partially redundant manner.

### Concentrations of polarity proteins do not correlate with cell size or bud number

Yeast cell polarity models display competition between polarity sites, on timescales that depend on the total amount of polarity proteins^19,25,41^. If cells allow a fixed time interval between polarity establishment and bud emergence, as appears to be the case in *S. cerevisiae*, then increased competition times could result in increasing numbers of buds^6^. Indeed, in cells of *S. cerevisiae* overexpression of polarity proteins can promote multi-polar outcomes or (in extreme cases) global Cdc42 activation^18,22,42^. In *A. pullulans*, multipolar outcomes are associated with increased cell size^34,36^. To determine whether cell size differences are associated with differences in polarity protein concentration, we compared fluorescence intensity of tagged proteins with cell volume. We converted fluorescence intensity to protein concentration using molecular brightness estimates from fluorescence correlation spectroscopy (FCS).

Measurements from tens of thousands of cells are summarized in Figure 2A and a more complete description of the experiments and data are in Figure S3. For all five polarity proteins, the average concentration varied little with respect to cell size (Figure 2E and Figure S3). We conclude that cells do not make more buds due to different concentrations of the core polarity factors. However, with similar concentrations of polarity proteins, larger *A. pullulans* cells would have increased total amounts of polarity proteins, which might explain why they can make more buds.

**Figure 2.**
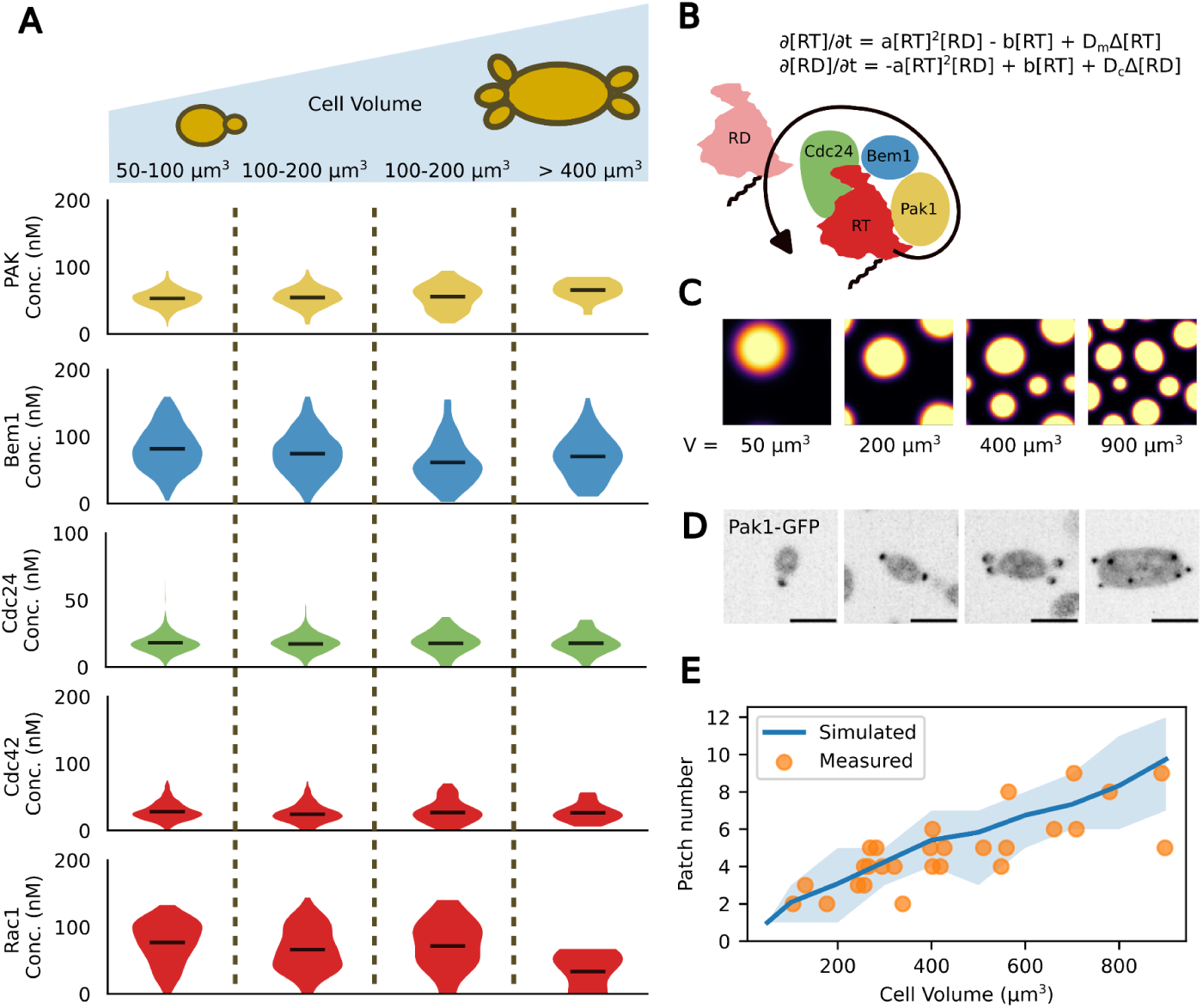
Polarity patch number increases with cell size. Polarity protein concentrations are similar regardless of cell size. (A) Cytosolic concentrations of each polarity protein were measured in live cells and binned according to cell size. The median brightness was measured in confocal images masked around individual cells and converted to concentration using the molecular brightness of the probes, which were determined using fluorescence correlation spectroscopy (FCS). (B) Minimal model of cell polarity. The PAK (Pak1), scaffold (Bem1), and GEF (Cdc24) interact with GTP-bound Rac1/Cdc42 (RT) to promote activation and membrane association of the inactive, GDP-bound Rac1/Cdc42 (RD). Only RT and RD are represented explicitly in the equations. Cooperativity is incorporated by making the rate of activation proportional to RT^2^. (C) Squares show results for the RT concentration of cells with different volumes. Yellow indicates regions of high RT concentration. Simulated domains were seeded with noisy initial protein concentrations and simulations were curtailed at 30 minutes. Total concentration of GTPase was the same across simulations. (D) Images of *A. pullulans* cells expressing Pak1-GFP. Larger cells appear to have more polarized sites at which buds are seen emerging. Scale bar, 5 μm. (E) Polarity patch number by cell size in simulations (blue line) and cells (orange circles). Simulations were performed with cell volumes of 50, 100, 200, 300, 400, 500, 600, 700, 800, and 900 μm^3^ and repeated 10 times for each volume with different random initial conditions (range of replicates is lighter blue ribbon).

To assess the plausibility of this hypothesis, we compared cell volume and polarity site number in cells and in a minimal model of cell polarity with only two molecular species, representing active and inactive forms of the Cdc42/Rac1 GTPase (Figure 2B). This minimal model is simpler to implement and optimize, but preserves several core features of a mechanistic polarity model developed for the *S. cerevisiae* polarity circuit^14^. We simulated the localization of the GTPase on a square domain with periodic boundaries while varying the cell volume and surface area as appropriate for a sphere. The concentration of GTPase in the entire system was assumed constant regardless of cell size, and simulations were initiated with heterogeneous concentrations of active GTPase using gaussian noise. When polarization was simulated for 30 minutes, the approximate minimum time between cell birth and bud emergence, the number of polarity sites increased with the size of the simulated cell (Figure 2C). Comparing our simulation results to the number of Pak1-GFP sites in *A. pullulans* cells revealed an excellent match (Figure 2D, E). Thus, a minimal model of cell polarity in which competition is cut off at 30 minutes quantitatively captures the relationship between cell volume and polarity site number in *A. pullulans*.

### Dynamics of polarity establishment in *A. pullulans* indicate local negative feedback

To assess whether curtailed competition is responsible for multipolarity in *A. pullulans*, we next followed the dynamics of polarity establishment using Pak1-GFP, the brightest of our polarity probes. Pak1 accumulated rapidly at polarity sites, and often transiently polarized at more sites than were ultimately maintained by the cell, consistent with competition between polarity sites (Figure 3A, 3B). The transient polarity sites disappeared before bud emergence, which generally occurred 15 to 30 minutes after initial polarization. Stable polarity patches also fluctuated in intensity, seeming to appear, disappear, and reappear in the same place over time. To analyze these dynamics we initially focused on small cells that formed only one bud. The Pak1 concentration at polarity sites often appeared to oscillate with a consistent period (Figure 3C-H; Figure S4). Intensity power spectra often showed a peak corresponding to a period of one or two minutes (Figure 3E), but in some cases were difficult to interpret due to the high weight of lower frequency signals attributed to random fluctuations in concentration via a Wiener process. Autocorrelation analysis of signal intensity was also suggestive of oscillations, with peaks indicating time lags at which the signal became self-similar again (Figure 3F). Using the first peak in the autocorrelation function to indicate the oscillation frequency, most polarity sites displayed a period of one to two minutes, with longer periods in some cells (Figure 3G). Oscillatory dynamics have been described in many biochemical systems^43,44^, and generally indicate the presence of negative feedback.

**Figure 3.**
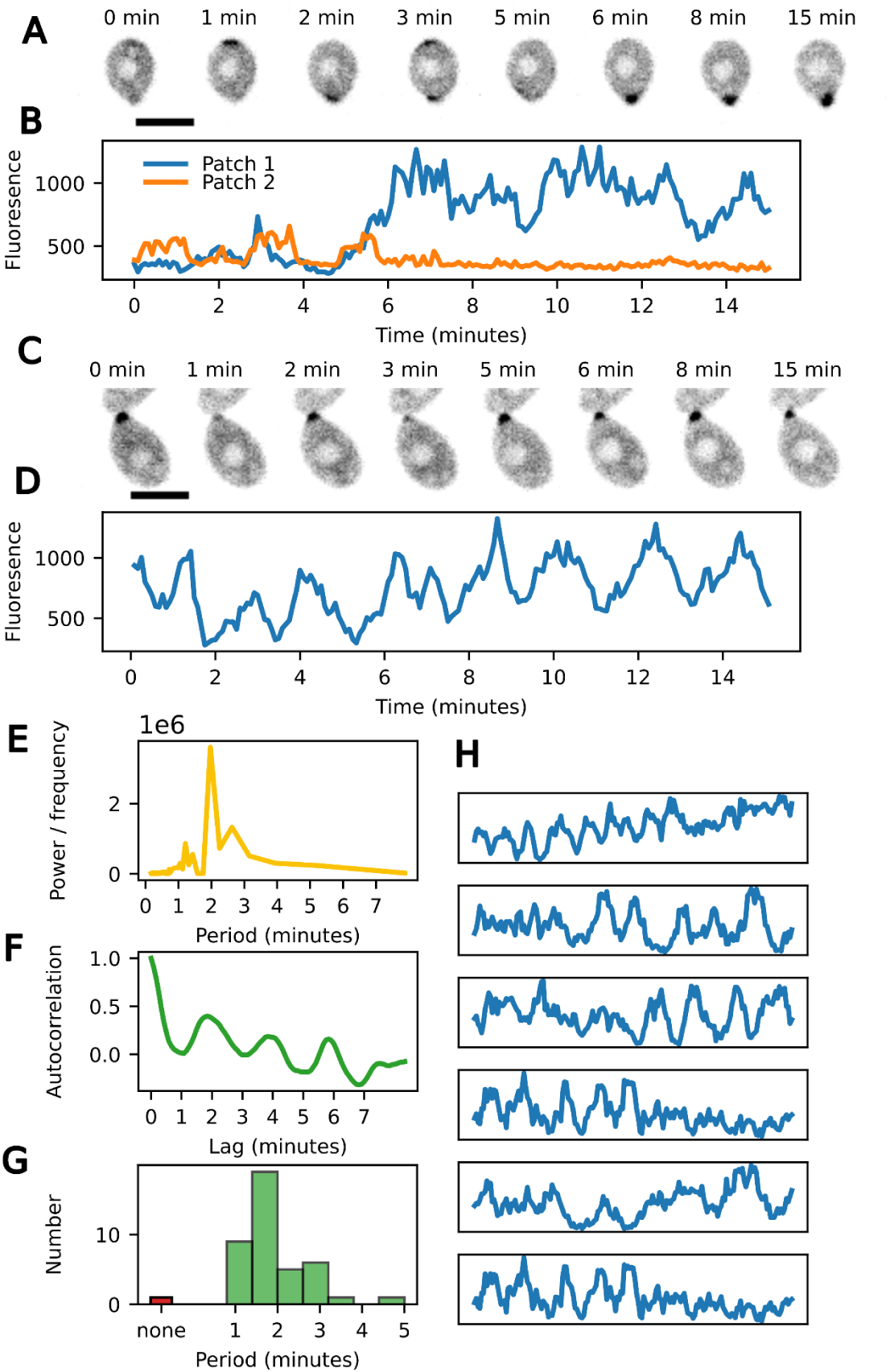
Dynamics of cell polarity in *A. pullulans* cells making one bud. (A) A cell polarizes transiently at two different sites. One site disappears and the other remains at the site of bud emergence. Scale bar, 5 μm (B) Maximum intensity traces of the polarity patches from A over time. (C) A single polarity site displays fluctuating intensity. Scale bar, 5 μm (D) Maximum intensity trace of the polarity patch from C over time. (E) Power spectral density of the signal from D. The peak indicates that the signal oscillates with a period of two minutes. (F) The autocorrelation function of the signal from D. The peaks correspond to lag times at which the signal becomes self-similar, consistent with a 2 min period. (G) Histogram showing the first peak in the autocorrelation functions of polarity signals from many cells with single polarity sites. A window of 3 neighbors on each side was used in calling peaks to filter out noise. (H) Example traces of polarity signals from more cells, highlighting the variability of these signals from cell to cell.

We next examined cells with more than one polarity site, to assess whether oscillations were coupled or independent of each other. To reduce the complexity of the analysis, we focused on cells with exactly two polarity sites. Polarity site intensity oscillated with similar frequencies in cells with two polarity sites and cells with only one (Figure 4A, 4B, 4C). There were example cells where two sites oscillated in phase or out of phase with each other, but there was no discernible pattern of coordination between sites in the same cell, and the same two sites could (through small differences in their periods) transition from in-phase to out-of-phase oscillations. To quantitatively measure any relationship between oscillations in the same cell, we calculated the crosscorrelation between site intensities. Any two oscillators with similar frequency should have peaks in their cross-correlation as well as their individual autocorrelations (Figure 4D), whether coupled or not. When averaging cross-correlations from many cell measurements, peaks with time-shifts that are consistent across cells should emerge. Instead, we observed an essentially flat cross-correlation when averaging many oscillator pairs (Figure 4E). The lack of any consistent cross-correlation suggests that individual polarity sites oscillate independently, which has implications for the mechanism of negative feedback (see below).

**Figure 4.**
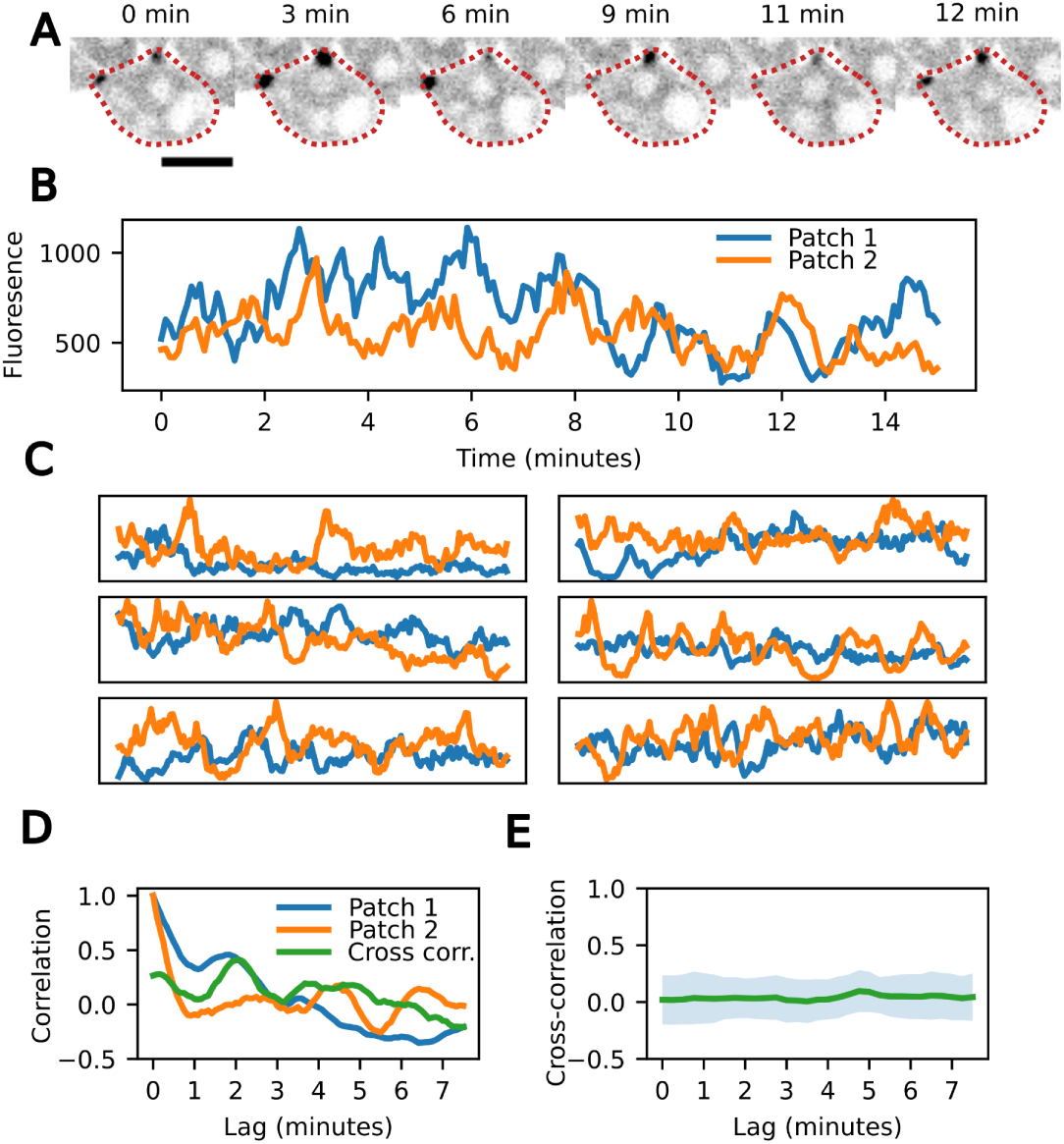
Dynamics of cell polarity in *A. pullulans* cells making two buds. (A) A cell polarizes two different sites. Each site oscillates over time. (B) Maximum intensity traces of the polarity patches from A over time. (C) Example traces of polarity signals from more cells. Signals from two patches in the same cell are similar in intensity to one another. (D) Autocorrelations of signals from each patch in A and the cross-correlation between them. Cross correlation shows peaks near zero and two minutes, suggesting nearly synchronous oscillations in this cell. (E) Average of cross-correlations of pairs of patches shows no peak, suggesting oscillations at each patch show no consistent time lag between them.

### Negative Feedback may serve to equalize polarity sites in multi-budded cells

Biochemical oscillations are indicative of negative feedback^43^, and negative feedback is also known to be one way to equalize polarity sites for steady-state multi-polar outcomes when incorporated into models of cell polarity^22,24,25,45^. To ask whether adding a negative feedback loop that promotes oscillations like those seen in A. pullulans would also promote equalization between polarity sites, we turned again to a computational model.

Negative feedback in both *S. cerevisiae*^30^ and *S. pombe*^46,47^ is thought to involve PAK-mediated inactivation of the GEF. While the mechanism of negative feedback in *A. pullulans* is unknown, we used GEF inactivation as the basis of negative feedback for our own model. In our model, active GTPase recruits both Cdc24 and Pak1 from the cytosol. Cdc24 promotes the activation and membrane association of the GTPase, providing positive feedback, while Pak1 promotes the return of Cdc24 to the cytosol, providing negative feedback (Figure 5A). With appropriate parameter choices, this model polarized to a single site in small cells and multiple sites in larger ones, and all polarity sites displayed oscillations (Figure 5B-D). When multiple polarity sites were maintained, their time-averaged intensities were very similar, indicating that the same model parameters can lead to both oscillation and equalization of polarity sites.

**Figure 5.**
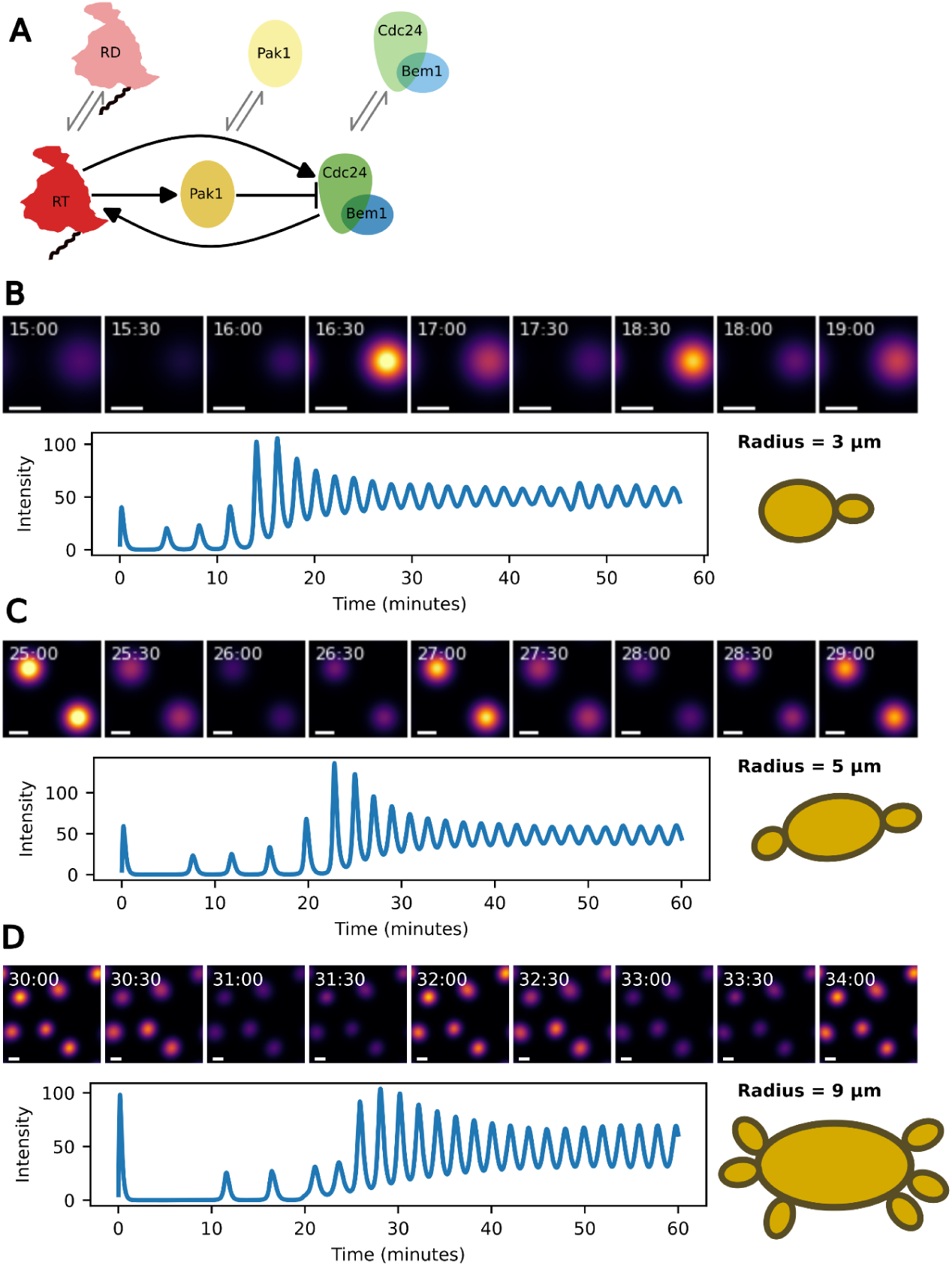
A polarity model incorporating negative feedback generates oscillating, and equalizing, polarity sites. (A) Schematic of the model architecture. Species in the top row represent the fast-diffusing cytosolic pool of each protein, while those in the bottom row represent the slow-diffusing membrane pools. Active, GTP-bound Rac1/Cdc42 (RT) at the membrane recruits both Pak1 and Cdc24 to the membrane. At the membrane, Cdc24 promotes the conversion of GDP-bound, cytosolic Rac1/Cdc42 (RD) into RT, constituting a 2-node positive feedback loop. At the membrane, Pak1 promotes the return of Cdc24 to the cytosol, constituting a 3-node negative feedback loop. (B) Simulation of a cell with a 3 μm radius. Top row shows RT concentration on the membrane over time. Scale bar is 1 μm. Lower row shows the maximum intensity of the single patch over the course of 1 h. (C) Simulation of a cell with a 5 μm radius. Top row shows RT concentration on the membrane over time. Scale bar is 1 μm. Lower row shows the maximum intensity of one of the two patches over time. Patches oscillate synchronously in this simulation. (D) Simulation of a cell with a 9 μm radius. Top row shows RT concentration on the membrane over time. Scale bar is 1 μm. Lower row shows the maximum intensity of one of the seven patches over time. Patches form at different times but eventually synchronize.

Unlike the oscillations we observed in *A. pullulans* cells (Figure 4), oscillations in model simulations were perfectly in-phase (Figure 5). Small changes to model parameters, however, resulted in anti-phase oscillations (Figure S5A). Although initialized with noisy distributions representing random molecular noise, our simulations were subsequently deterministic, meaning that even weak coupling effects are sufficient to synchronize oscillations.

To understand whether the equalization emerges from negative feedback or some other feature of the model, we constructed a similarly parameterized model lacking PAK-mediated negative feedback (Figure S5B). In this model, no oscillations were present, as expected. As with our previous minimalistic model (Figure 2B,C), a single polarity patch developed when small domains were simulated, while multiple sites could be maintained with larger simulated domains (Figure S5C-E). When multiple sites were present, they competed, and competition was slowed for very large domains.

### Pak1 provides negative feedback and promotes equalization of polarity sites

PAK-mediated GEF inactivation is one way that negative feedback can be incorporated into the polarity circuit^30,46^ but alternative mechanisms have also been proposed^22,25,31,48,49^. To evaluate whether Pak1 contributes to the negative feedback we observed in *A. pullulans*, we monitored cell polarity dynamics in a *pak1*Δ mutant. Since we could not use Pak1-GFP to monitor polarity in the *pak1*Δ cells, we used a Bem1 probe to monitor polarity. Because the Bem1-GFP signal was weak relative to Pak1-GFP, we used a strain in which a 3xGFP-BEM1 reporter is expressed from a heterologous (*S. cerevisiae ACT1*) promoter. To validate this probe, we constructed a strain expressing both Pak1-mCherry and 3XGFP-Bem1. The two probes were colocalized, and displayed simultaneous fluctuations in intensity (Figure S6).

The 3XGFP-Bem1 probe polarized to distinct sites in both the wildtype and *pak1*Δ mutant strains (Figure 6A, 6F), but showed diminished fluctuations in fluorescence intensity in the mutant (Figure 6B, 6C, 6G, 6H). While autocorrelation of polarized signals in the wildtype showed clear peaks (Figure 6D), autocorrelation functions in the *pak1*Δ did not (Figure 6I). We identified the first peaks in the autocorrelation functions by finding their first local maxima, using a window of three neighbors on each side to smooth maxima so that small fluctuations as seen in Figure 6I were not spuriously identified as peaks. While the peaks in a wildtype strain were mostly between one and two minutes (Figure 6D), peaks in the *pak1*Δ cells occurred at longer, less consistent time delays or often not at all (Figure 6J, Figure S7). We conclude that Pak1 is important for generating oscillations in polarity site intensity, consistent with a role for Pak1 in providing negative feedback in *A. pullulans*.

**Figure 6.**
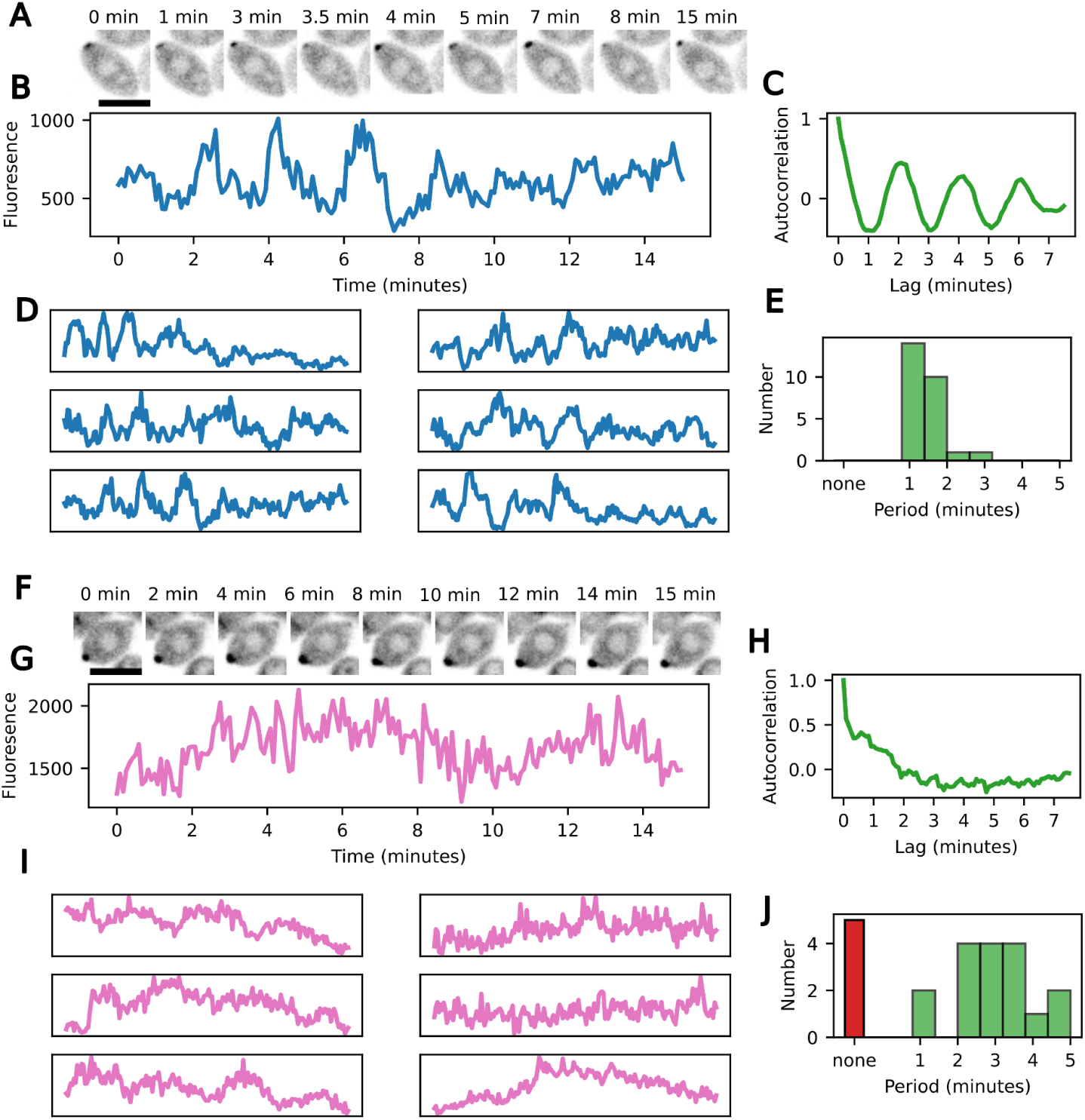
Cells lacking *PAK1* polarize, but polarity sites do not oscillate. (A) Example wildtype cell with one polarity site visualized with 3xGFP-Bem1 probe. Scale bar, 5 μm. (B) Maximum intensity over time for the polarity site shown in A, showing oscillation. (C) Autocorrelation of signal from B. The first peak at about two min indicates oscillation period. (D) Additional examples of maximum intensity traces from wildtype cells as in B. (E) Histogram showing the first peak in the autocorrelation functions of polarity signals from many cells with single polarity sites. A window of 3 neighbors on each side was used in calling peaks to filter out noise. (F) Example pak1Δ cell with one patch of polarized 3xBem1-GFP probe. The patch persists over time, leading to bud emergence in the final frames. Scale bar 5 μm. (G) Maximum intensity over time of the polarity patch from the pak1Δ cell in F. Fluctuations at very short time scales are apparent but oscillations every one-to-two minutes are not. (H) Autocorrelation of the signal appears to simply decay with increasing lag time, as expected for a simple brownian noise process. (I) Additional examples of maximum intensity traces from pak1Δ cells with one polarity patch. (J) Histogram showing the first peak in the autocorrelation functions of polarity signals from many pak1Δ cells with single polarity sites. A window of 3 neighbors on each side was used in calling peaks to filter out noise.

As computational modeling suggested that negative feedback in the polarity circuit could equalize polarity patches, we tested whether Pak1-mediated negative feedback might serve to equalize polarity patches in cells. Because instantaneous intensity oscillates over time in wildtype cells, we measured the average intensity of polarity sites over 15 minutes between initial polarization and bud emergence (Figure 7A, 7B). We observed that the differences in intensity between two patches from the same cell were much smaller in the wildtype than the *pak1*Δ mutant, indicating that Pak1 is required for patch equalization (Figure 7C). Moreover, *pak1*Δ mutants cells often accumulated the Bem1 probe to higher levels than those in wild type cells, consistent with failure to constrain patch growth through negative feedback.

**Figure 7.**
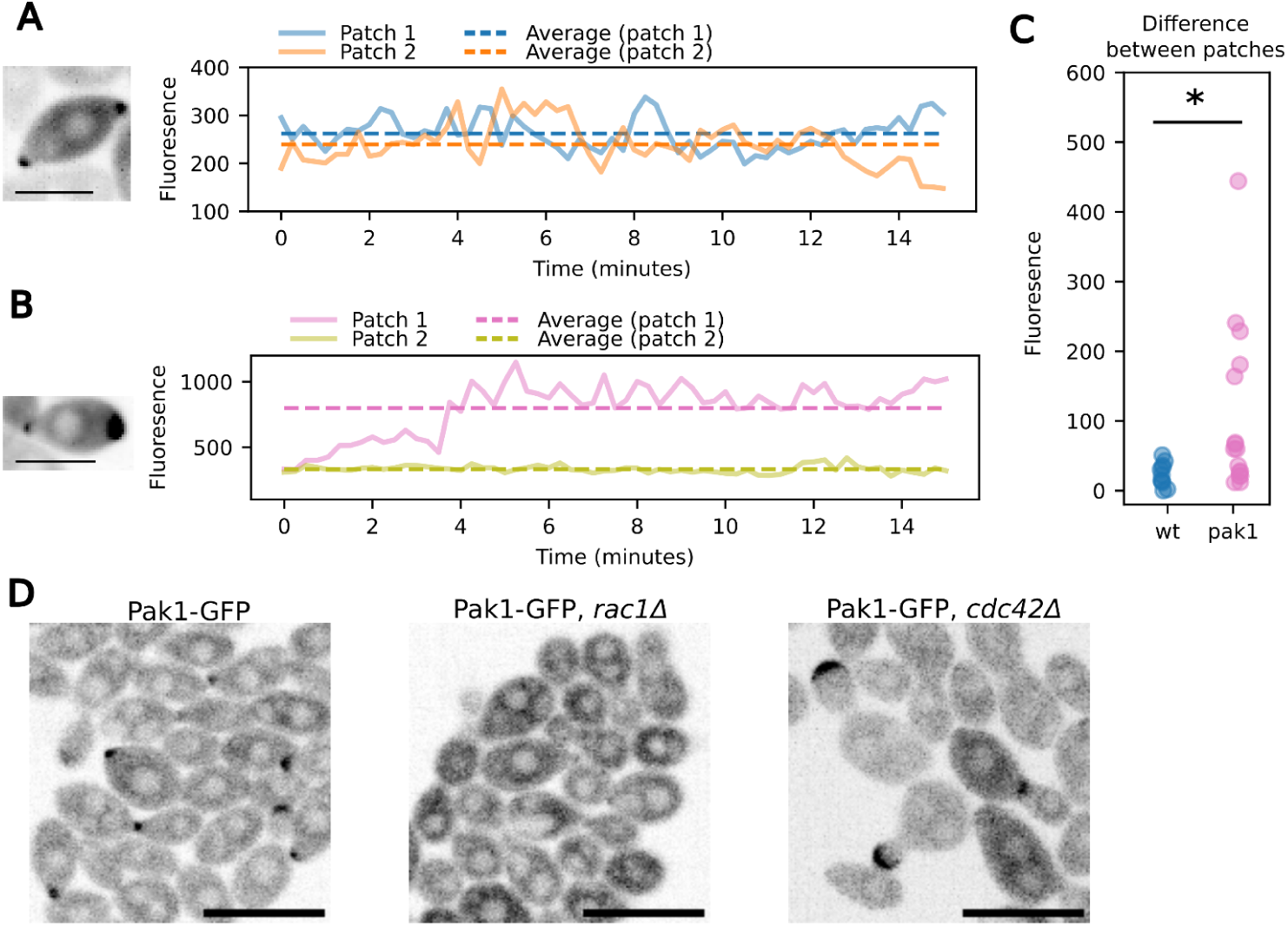
Cells lacking PAK1 fail to equalize polarity patches. (A) Example wildtype cell with two patches visualized with 3xGFP-Bem1 probe. Left: mean intensity projection of images taken every 15 seconds for 15 minutes in order to average the fluctuations in intensity. Scale bar, 5 μm. Right: maximum intensity of each patch over time. Dashed lines indicate averages over the entire interval. (B) Example *pak1*Δ cell with two patches, analyzed as in A. (C) Intensity differences between two patches in the same cell. Differences are consistently small in wildtype but larger and more variable in *pak1*Δ cells (Mann-Whitney U test; p < 0.05). (D) Polarization of Pak1-GFP is very weak in *rac1*Δ cells, but remains strong in *cdc42*Δ cells.

PAKs directly bind and are activated by Rac/Cdc42-family GTPases^13,50–53^. As there are two such GTPases in *A. pullulans*, we tested whether Pak1 polarization was dependent on Cdc42 or Rac1. To that end, we deleted either *CDC42* or *RAC1* in a strain expressing Pak1-GFP. Localization of Pak1-GFP in the *cdc42*Δ mutant was similar to that in wildtype cells, while the probe was predominantly cytoplasmic in the *rac1*Δ mutant (Figure 7D). Thus, Rac1 is presumably the primary GTPase that activates Pak1 in *A. pullulans*.

## DISCUSSION

Cell polarization is a fundamental and widely conserved aspect of cell physiology, which has been adapted to regulate diverse morphological processes^11,54,55^. Mechanistic aspects of cell polarity circuits are best understood in the model yeasts *S. cerevisiae*, where a core polarity circuit reliably produces a single polarity site, and *S. pombe*, where tip factors external to the same polarity circuit reliably produce exactly two polarity sites at opposite ends of the cell^11,56–58^. While most budding and fission yeasts display similar patterns, a subset of yeasts including A. pullulans have developed a multibudding lifestyle in which different cells produce variable numbers of buds in each cell cycle^34^. For these and other multipolar cells, the mechanisms by which specific numbers of polarity sites develop remain unclear. Here, we present an initial characterization of the *A. pullulans* polarity circuit. Our findings are consistent with the idea that the core polarity circuit is similar to that in model yeasts, with both positive and negative feedback loops. Polarity site number increases with cell size, and a negative feedback loop plays an important role in ensuring that polarity sites accumulate equal amounts of polarity factors and produce similarly sized buds. These findings form the basis for understanding how multipolar cells develop a suitable number of polarity sites.

### The *A. pullulans* polarity circuit

*A. pullulans* expresses two polarity GTPases, Cdc42 and Rac1, that accumulate at polarity sites together with the GEF Cdc24. Cdc42 and Rac1 were individually dispensable for budding, while Cdc24 was essential. The simplest way to interpret these findings is that Cdc42 and Rac1 play partially redundant roles, and that as in *S. cerevisiae* and *S. pombe*, there is a positive feedback loop whereby local accumulation of GTPase promotes local accumulation of GEF, which activates more GTPase. However, whereas in the model yeasts this positive feedback loop is mediated by the scaffold Bem1, the sole apparent *A. pullulans* Bem1 was dispensable for budding (although the protein did accumulate at polarity sites). This suggests that the full mechanism(s) of positive feedback may differ between yeasts.

In *S. cerevisiae*, genes that are essential for budding are also essential for proliferation. However, in *A. pullulans*, *cdc24* mutants were able to proliferate (albeit slowly) without budding. Mutant cells grew isotropically to form large spheres that could subsequently undergo medial division. This alternate growth modality is consistent with the well-appreciated morphological plasticity of this organism, and may enable it to tolerate and thrive in multiple environments^32,33,59^.

Polarity sites in *A. pullulans* displayed oscillatory accumulation of polarity factors, indicative of a negative feedback loop in the polarity circuit. Oscillations occurred with somewhat variable periods of 1-4 min, and polarity sites in the same cell appeared to oscillate independently, suggesting that negative feedback acts locally rather than globally. Oscillations were largely absent in mutants lacking the GTPase effector kinase Pak1, suggesting that as in the model yeasts^30,46^, negative feedback is mediated by PAK-family kinases.

Although instantaneous levels of polarity factors differed at the polarity sites in multipolar cells, the time-averaged levels of factors at each site were remarkably consistent. This equality of polarity sites may ensure that different sites produce similar buds. In contrast, polarity sites could accumulate quite different levels of polarity factors in *pak1* mutants, and also in *rac1* mutants that largely failed to polarize Pak1. These findings suggested that Pak1, and perhaps the negative feedback mediated by Pak1, is important for polarity site equalization.

### A computational model reproduces several features of *A. pullulans* polarization

We developed a simplified computational model of cell polarity that incorporates both positive and negative feedback. The model includes three distinct protein species: a GTPase, a GEF, and a PAK (the activities of GAPs, GDIs, and phosphatases were included implicitly as first-order reactions). Each species can transition between an inactive form in the cytoplasm and an active form at the membrane. The GEF promotes nonlinear activation of the GTPase, which promotes local activation of the GEF in a positive feedback loop. The PAK is activated by the GTPase and promotes inactivation of the GEF, constituting a negative feedback loop. Because all active (membrane) species diffuse slowly, both feedback loops act locally. However, because the total amounts of each species are assumed constant, depletion of any cytoplasmic species would slow its activation globally. Parameters were chosen to reproduce the oscillatory accumulation of polarity factors at individual polarity sites that we observed in cells.

This simple model successfully captured several key features of polarization observed in *A. pullulans*. First, the model generated polarity patches that, once established, remained stable in space while oscillating over time. Second, as the size of the simulated cell increased, the model produced more polarity sites. Third, when multiple polarity patches were maintained, their time-averaged intensity was consistent across patches. This latter feature was due to the negative feedback loop, because a comparable model lacking the PAK developed polarity sites of different size and protein content. Thus, as in cells, the PAK promotes both oscillation and equalization, suggesting that negative feedback would suffice to explain equalization of polarity sites.

Although oscillations at different sites in the same cell are not coupled, oscillations in the model were perfectly in-phase. We suspect that this phase coupling emerges from the global depletion of cytoplasmic species, mentioned above. However, even in the model, the coupling was weak. By perturbing the system, we were able to induce anti-phase instead of in-phase oscillations. Our simulations were deterministic, lacking the molecular noise which might obscure coupling in a biological context. Alternatively, there may be additional complexity in the cell polarity circuit that we did not include in our model.

### Conclusion

We report the first analysis of the polarity circuit in a multibudding cell. Our findings suggest that core elements identified in model yeast polarity circuits, including positive and negative feedback loops, have been adapted in *A. pullulans* for its own morphological program. We suggest that, as in computational polarity models, increased cell size can enable the coexistence of more polarity sites, and that negative feedback can act to equalize the protein content of those polarity sites. Our findings also raise fascinating questions for the future. In particular, the precise mechanisms of positive and negative feedback remain to be determined, and the specific roles of Cdc42 and Rac1 also remain unclear. Further studies are likely to shed light on these questions.

## ACKNOWLEDGEMENTS

We thank the Gladfelter, Elston, and Lew labs along with Christian Zimmermann, Cene Gostinčar, and other members of the *A. pullulans* working group consortium for stimulating discussions and feedback on this manuscript. This work is supported by NSF Grant MCB2016022 to A.S.G.

## METHODS

### EXPERIMENTAL MODEL AND STUDY PARTICIPANT DETAILS

#### Yeast strains

Mutant strains of *Aureobasidium pullulans* were constructed in the EXF-150 genetic background. Gene deletions and fluorescent protein fusions were generated using homologous recombination by means of chemical transformation^36^. For each strain 3 separate transformants were checked for consistent phenotypes/localization to ensure that an individual transformant phenotype was not due to potential unrelated mutations. The GFP and mCherry tags were designed such that each amino acid was encoded with its most frequently used codon, based on all predicted coding sequences from the EXF-150 reference genome, as described previously^35^. Either hygromycin-or nourseothricin-resistance genes were incorporated into the DNA constructs for endogenous fluorescent tags and gene deletions. For fluorescently tagged proteins expressed as additional copies, probes were integrated at the *URA3* locus of a strain in which the *URA3* gene had been replaced with a hygromycin resistance gene^36^. A full list of the strains used in this study can be found in Supplemental Table 1.

#### Cell growth and imaging conditions

Cells were grown in 1% yeast extract, 2% bacto-peptone, 2% dextrose (YPD) medium at 25°C overnight before imaging. We applied 4 μL of culture to an Ibidi 8 well glass bottom μ-Slide and placed a pad of complete synthetic medium (CSM) with 2% agarose on top. Pads were punched from plates of agarose using a 1 mL pipette tip. We performed all imaging at room temperature.

### METHOD DETAILS

#### FCS and imaging

FCS measurements and concentration quantification were performed using a Zeiss 980 confocal microscope. For FCS calibration, one chamber of an Ibidi 8 well glass bottom μ-Slide was blocked with BSA, washed twice with water, and filled with dye. FCS calibration for green fluorophores was performed with Atto488. Initially, a dilution series of 16 μM, 8 μM, 4 μM, and 2 μM Atto488 was used to estimate focal volume and aberration in Z (FV = 0.42 μm^3^; k = 4.2). Subsequently, 2 μM Atto488 was used for calibration. Calibration for red fluorophores was performed with 2 μM Atto594 (FV = 0.43 μm^3^; k = 5.83). FCS measurements in cells were collected for 5 seconds, with one measurement per cell, and autocorrelations from a minimum of 10 cells were averaged for each probe. Confocal images were taken using a 40X water objective. A single position in Z was used, making each pixel in the resulting image the integrated fluorescence of a consistent volume.

#### Cell segmentation determination of protein concentration from fluorescence brightness

From still images of *A. pullulans* cells in a single Z plane, we generated masks around individual cells using the Cyto3 segmentation model in Cellpose3. Masks and images were imported into python, where tools from scikit-image were used to extract features from masked individual cell images. The long and short axes were extracted as features and used to estimate cell volumes as V = 4/3πr_long_r ^2^. The median pixel intensity value was used to estimate the cytosolic concentration within the cell. Median was chosen over mean to avoid enriched regions, due to polarization, and deficient regions, due to exclusion from vacuoles, lipid droplets, and nuclei. To convert pixel intensity to concentration, we used molecular brightness estimates obtained by FCS of Pak1-GFP, for green probes, or Rac1-mCherry, for red probes.

#### Time-lapse imaging

Live cell time-lapse imaging was performed on a Nikon spinning disk confocal microscope with a 40X silicon-immersion objective. When using Pak1-GFP expressing strains to monitor polarity patch dynamics in cells with only one patch, we took images every 5 seconds in a single Z position. To capture polarity dynamics at multiple sites in the same cell we collected multiple Z positions at 15 second intervals and analyzed maximum intensity projections. For cells with exactly two patches, we were able to use some 5 second interval/single Z movies in favorable circumstances, as well as 15 second interval/maximum intensity projections from 7-11 Z positions. Oscillatory behavior was similar with both acquisition protocols. To monitor polarity dynamics using the 3XGFP-Bem1 probe, Z stacks were taken at 15 second intervals.

### QUANTIFICATION AND STATISTICAL ANALYSIS

#### Analysis of polarity patch dynamics

To analyze the dynamics of polarity patches, we cropped time-lapse images around individual cells before loading and processing the cropped movies using custom scripts written in the Julia programming language^60^. In short, projections were made from each image such that each pixel represented the standard deviation of the fluorescence at that position with respect to time. We found that this method gave much stronger separation of cytoplasmic signal and polarized signal than did a more intuitive maximum intensity projection image, and produced high signal at the entire area traversed by the polarity patch, making thresholding robust to small amounts of cell movement during the time-lapses. From the projections, we thresholded at half of the maximum signal to find the polarity sites. For each polarity site, we then took the maximum intensity of the site at each time point in the time-lapse movie. The movies were 30 minutes, but the signals were then cropped to 15 minutes, beginning at the time at which the signal first reached half its maximum. These time-cropped signals were used to generate the autocorrelations and the periodograms of these signals. The code used to generate these figures can be found in the GitHub page associated with this article.

#### Analysis of patch number as a function of cell size

A model of cell polarity, which included only two molecular species, was based on equations described previously^14,19^, and manually re-parameterized so that patch number scaled with cell size similarly to measurements in *A. pullulans*. The cell membrane was simulated as a square grid, and the volume of the simulated cell was calculated as if the surface area of that grid corresponded to a sphere. The total amount of GTPase in the system was scaled so that its concentration remained constant. The grid was discretized as 100 x 100 points, and diffusion of species was adjusted accordingly. To construct random initial conditions, RT concentrations on the membrane include Gaussian noise. Details for the numerical scheme used to simulate the equations is provided below.

Polarization was simulated for 30 minutes, and the number of patches at that time was determined by finding local maxima with intensity values greater than 10 nM. Equations are written below and parameters can be found in the supplement and in the code repository associated with this publication. The parameter *η* is a conversion factor for concentrations at the membrane to concentrations in the cytosol^14^ and depends on the size of the cell that is simulated.

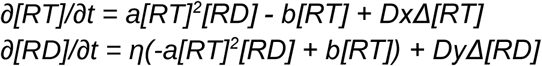

For measurements in live cells, we manually counted the number of polarity patches in budding cells, and approximated the volume of the corresponding mother cells by assuming it was an ellipsoid of volume V = 4/3πr_long_r_short_^2^, where width is the shorter axis. The images were maximum intensity projections of Pak1-GFP cells grown at low densities, a condition which favors large, multi-budding cells.

#### Modelling polarity with negative feedback

For our model which included negative feedback, the reaction network was formulated to be as minimal as possible: We did not include the explicit formation of any protein complexes, and only three membrane-associated species were used. The recruitment of GTPase to the membrane was the product of the cytosolic concentration of the GTPase and the square of the concentration of membrane-associated GEF. Using the square of the GEF concentration ensured sufficient nonlinearity of the positive feedback, without necessitating additional species complexes, and is consistent with reports that *S. cerevisiae* Cdc24 can dimerize^61^. The membrane-associated PAK and GEF were each recruited as the product of their cytosolic concentrations and the concentration of membrane-associated GTPase. PAK accumulated more slowly and promoted the membrane-dissociation of the GEF as a product of the membrane-associated GEF and PAK concentrations. Equations for this model are:

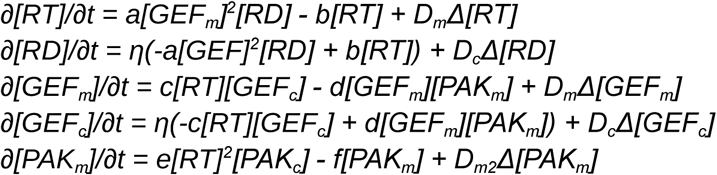

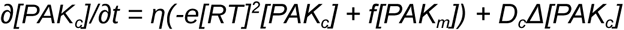

The results of a similar model, but lacking PAK-mediated negative feedback, are included in Figure S4. The equations for that model are:

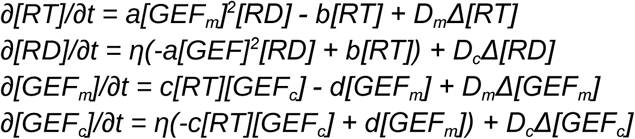

Parameters for each model can be found in Table S4 and Table S5, as well as the code repository associated with this publication.

#### Numerical Method

PDEs were solved numerically using a finite difference method in the Julia programming language^60,62^. Spatial domains were discretized as 100x100 square grids with periodic boundary conditions; timesteps were automatically determined per-simulation by the adaptive-time solver, CVODE, which performed time integration steps using the generalized minimal residual method (GMRES)^63–65^. Absolute and relative error tolerances were required to be at most 10^-8^ and 10^-5^ respectively to ensure numerical stability.

## ADDITIONAL RESOURCES

Code used to analyze data and generate figures is available at our GitHub repository: https://github.com/Gladfelter-Lab/Crocker2025/

## SUPPLEMENTAL INFORMATION

Document S1. Figures S1-S8 and Tables S1-S5

