## Supplementary material for "Negative feedback equalizes polarity sites in a multi-budding yeast": Document S1. Figures S1-S8 and Tables S1-S5

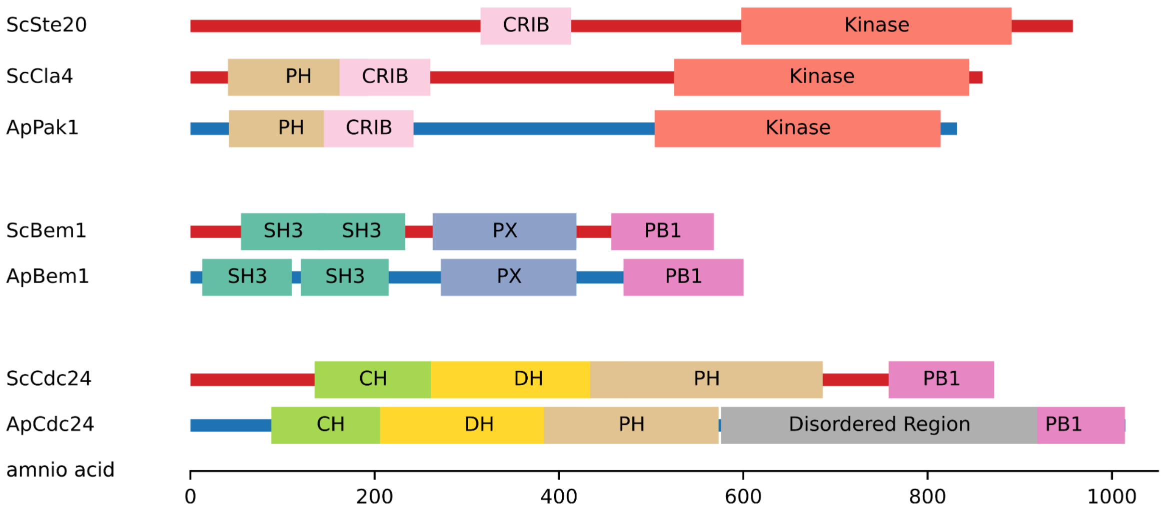


**Figure S1. Domain comparison of *Saccharomyces cerevisiae* polarity proteins and their homologs in *Aureobasidium pullulans*. Related to Figure 1.**

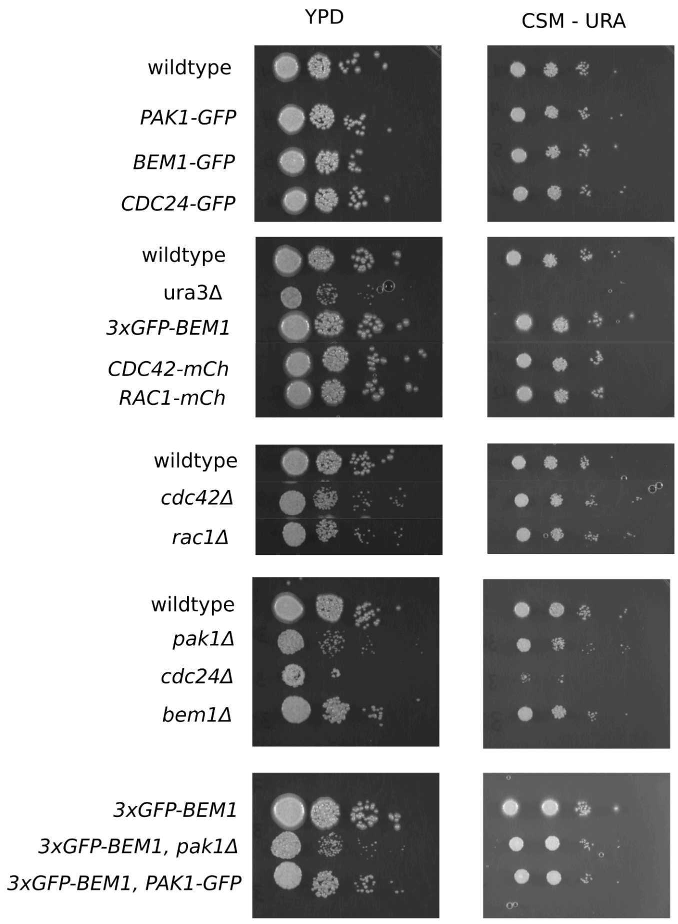


**Figure S2. Spot plates of each strain used in this study.**

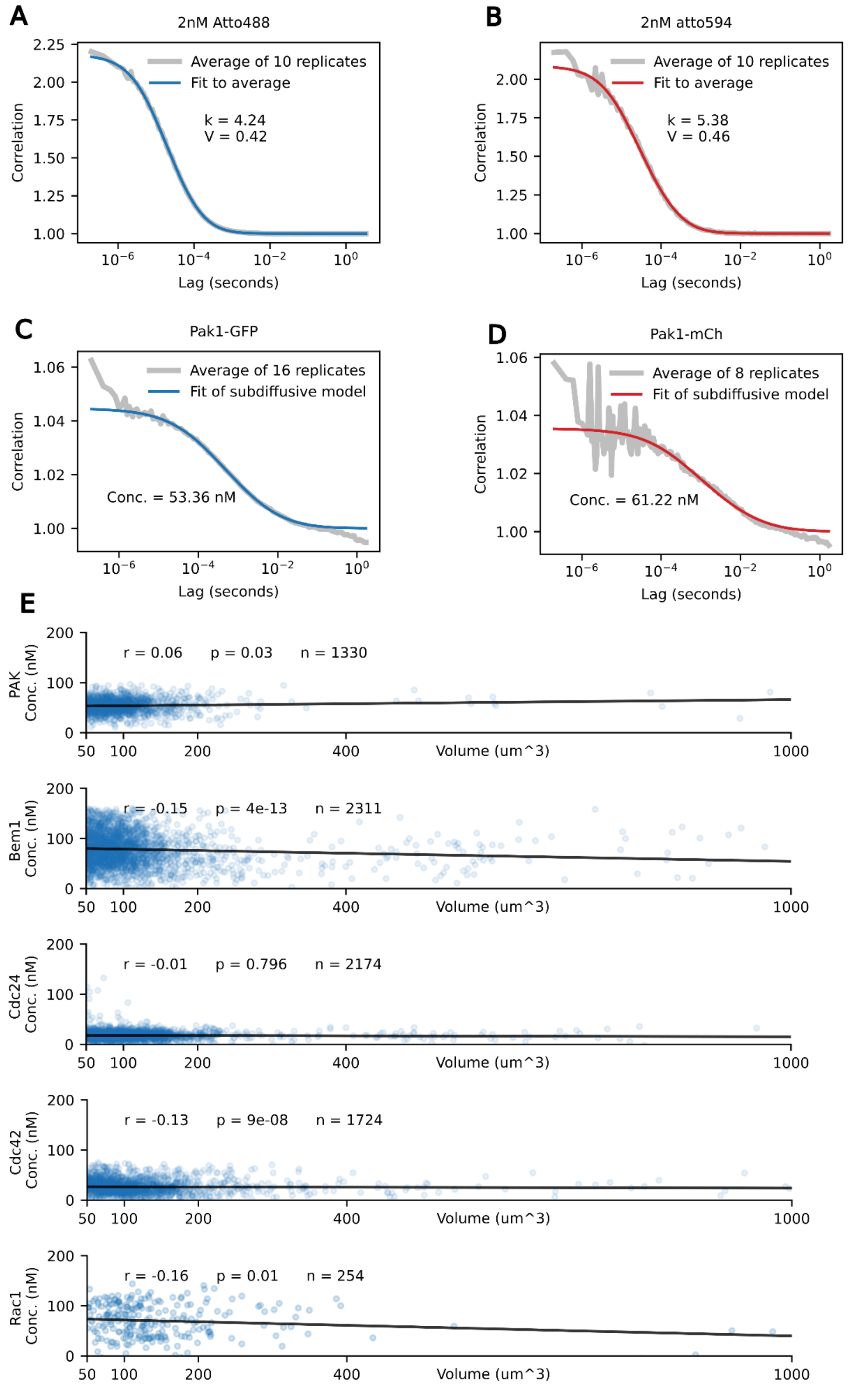


**Figure S3. (A-D) FCS calibration and measurement. (E) Full concentrations and size data associated with Figure 2.**

A

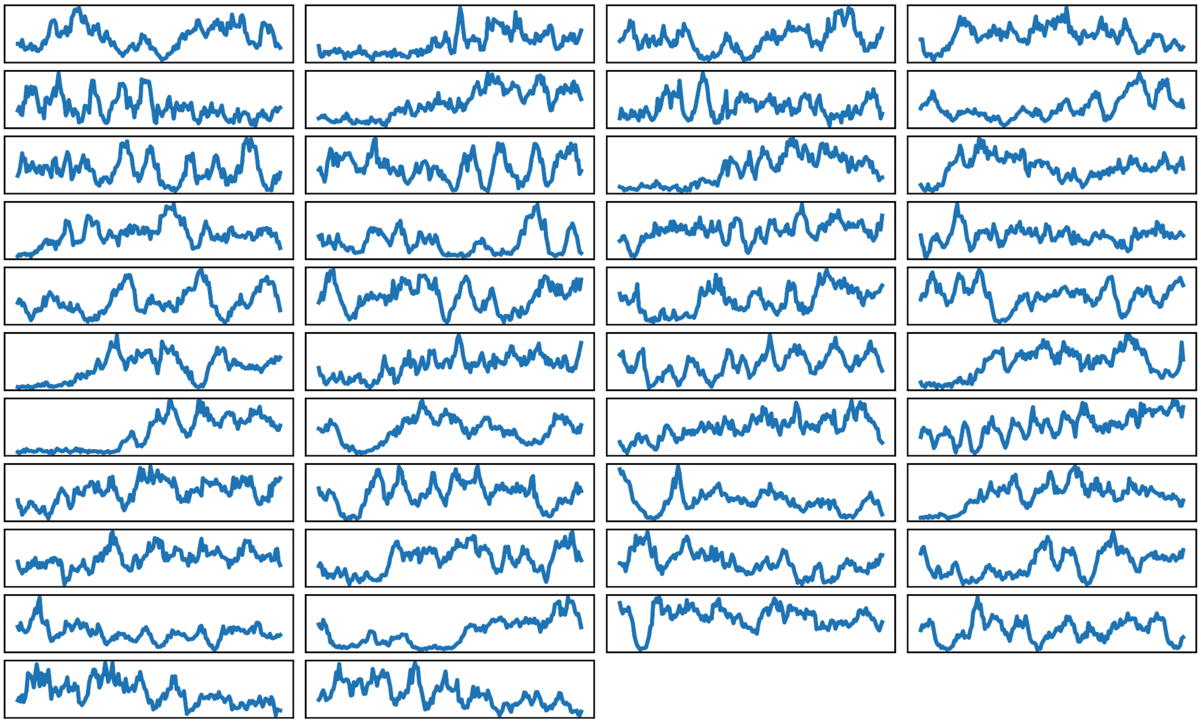

B

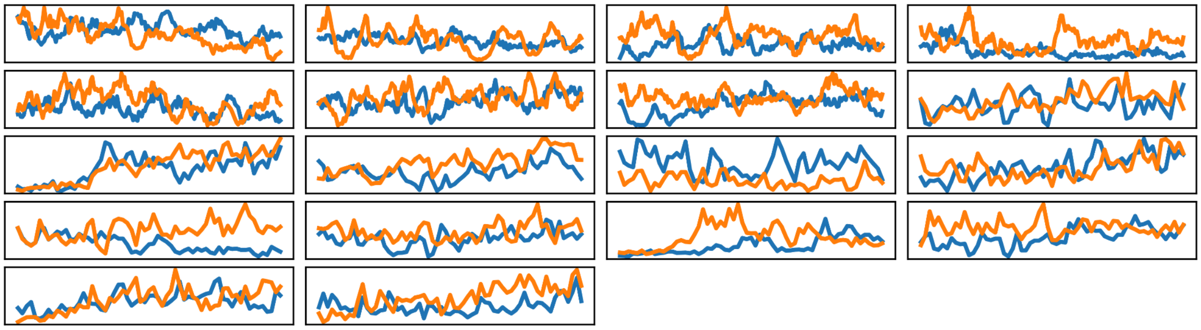

**Figure S4. Traces from all polarity patches of all cells. Related to Figure 5 and Figure 6.**

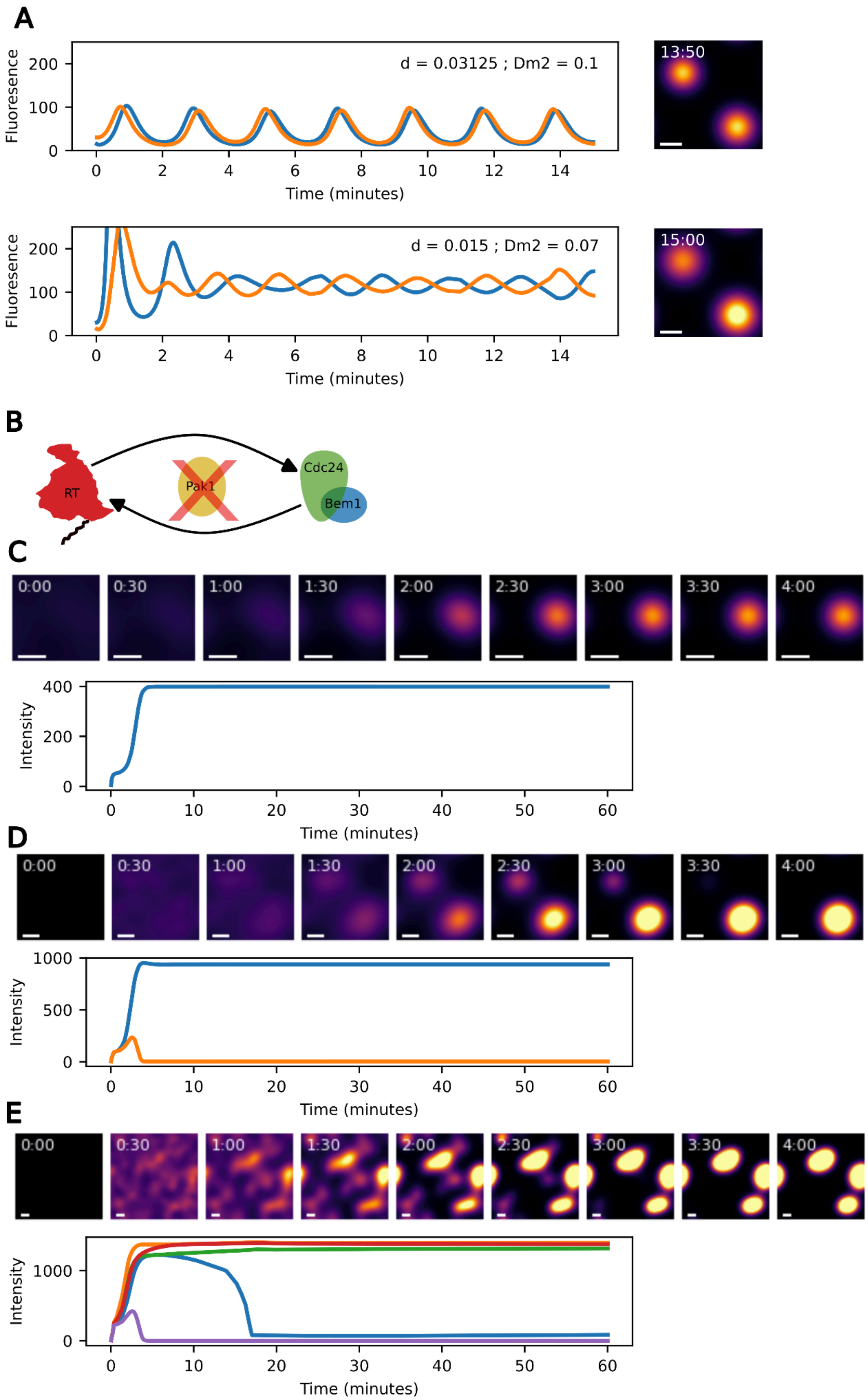
A
d = 0.03125 ; Dm2 = 0.1
13:50
Fluoresence
Time (minutes)
d = 0.015 ; Dm2 = 0.07
15:00
B
RT
Pak1
Cdc24
Bem1
C
0:00
0:30
1:00
1:30
2:00
2:30
3:00
3:30
4:00
Intensity
Time (minutes)
D
Intensity
Time (minutes)
E
Intensity
Time (minutes)

**Figure S5. (A) Coupling of polarity patches is sensitive to parameterization. After reaching synchronous oscillations as in Figure 5, the concentration of each species on the membrane was halved for one site but not the other. Simulations show the response over 15 minutes after that perturbation. The initial parameterization returns to oscillating in-phase; an alternative parameterization oscillates out-of-phase. (B-E) When negative feedback is removed from the polarity model from Figure 5 sites compete, do not oscillate, and accumulate higher levels of active GTPase (RT).**

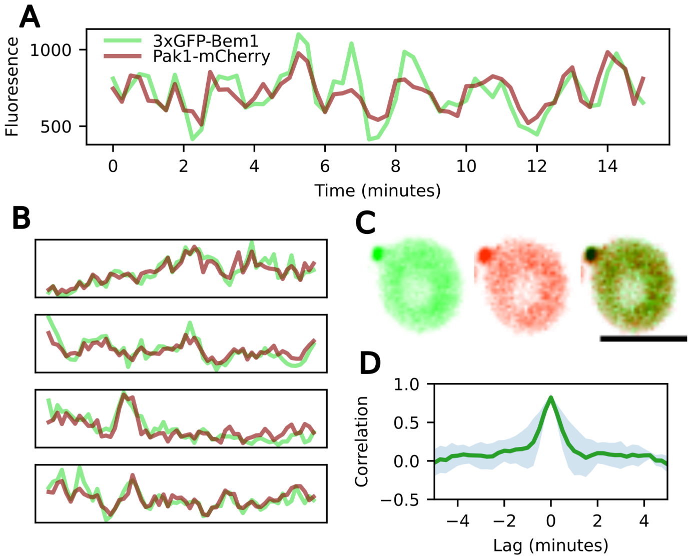


**Figure S6. Bem1 and Pak1 probes colocalize and covary without a detectable delay. Related to Figure 6.**

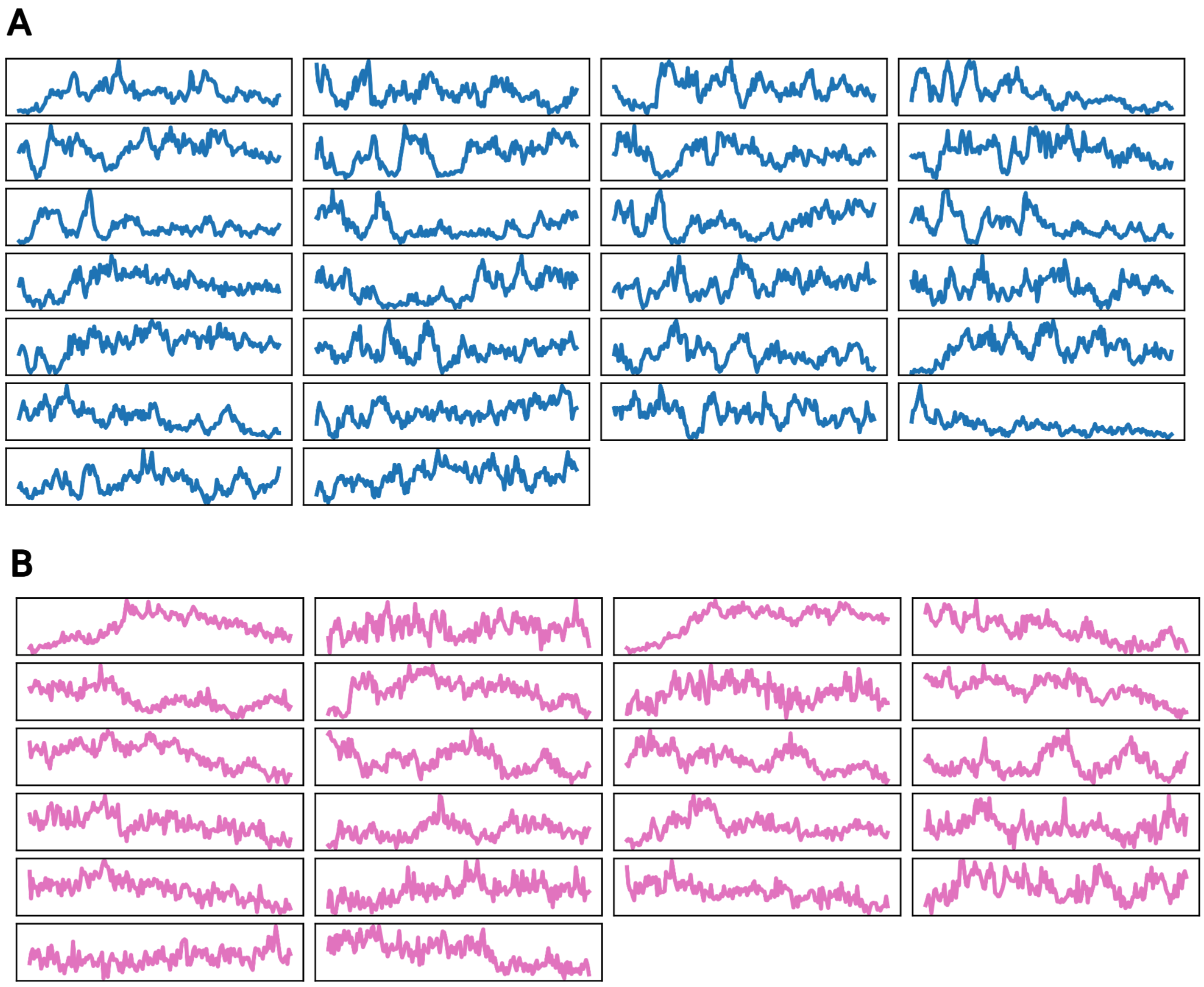


**Figure S7. Traces from each single budded cell with *PAK1* (A) and each *pak1Δ* (B). Related to Figure 6.**

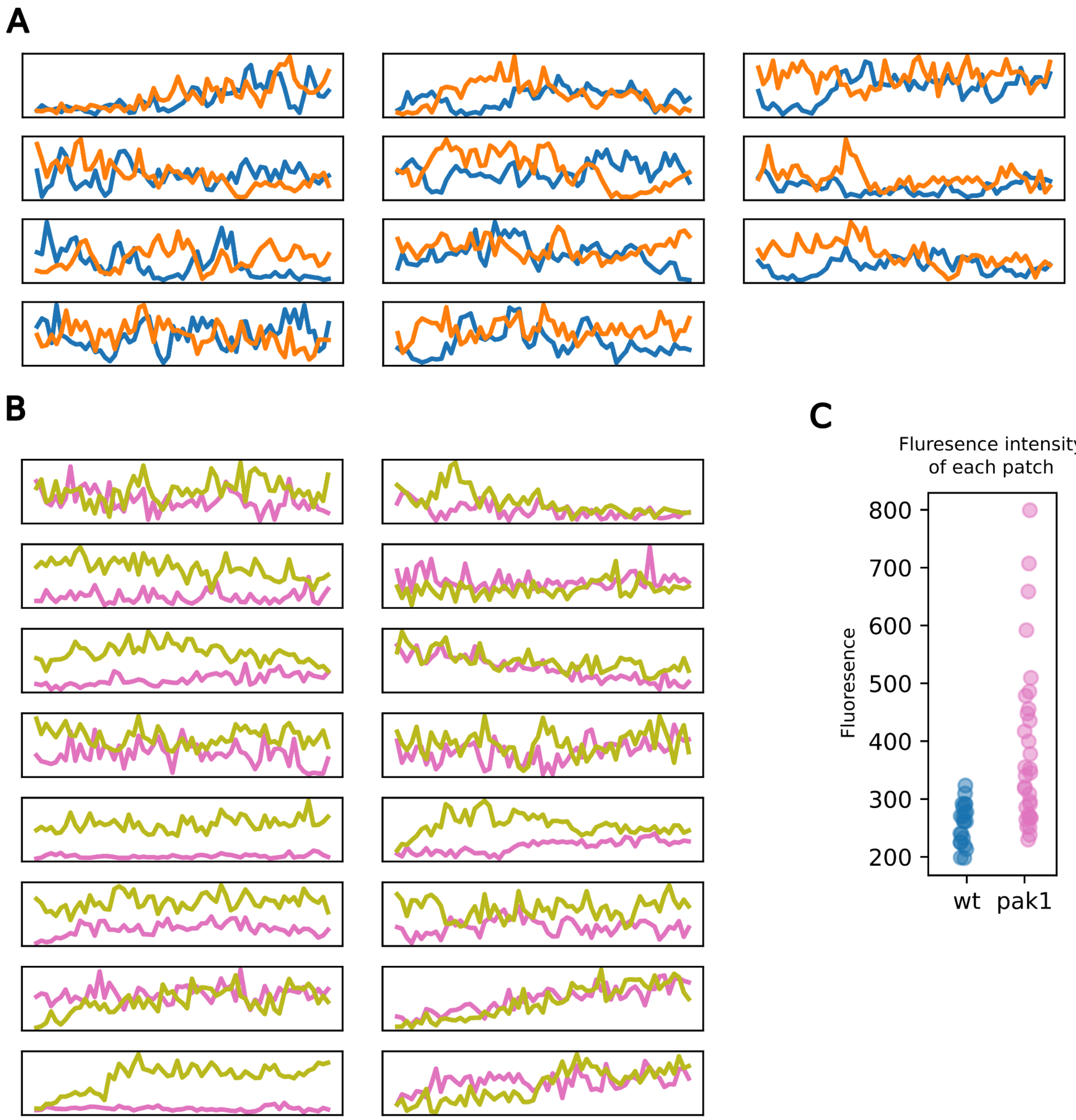


**Figure S8. Traces from each two-budded cell with *PAK1* (A) and each *pak1Δ* (B). Difference between each set of polarity patches (C). Related to Figure 6.**

| Protein Name | NCBI Accession | *S.c.* Homolog | Query Cover | E value | Per. Ident | RBH |
|---|---|---|---|---|---|---|
| Pak1 | XP_029760918 | Ste20 | 49% | 1.00E-93 | 46.90% | Yes |
| Bem1 | XP_029763853 | Bem1 | 70% | 9.00E-54 | 33.70% | Yes |
| Cdc24 | XP_029763714 | Cdc24 | 52% | 2.00E-40 | 27.16% | Yes |
| Cdc42 | XP_029765618 | Cdc42 | 98% | 2.00E-113 | 78.39% | Yes |
| Rac1 | XP_029765527 | Cdc42 | 99% | 6.00E-95 | 67.19% | No |
| Rdi1 | XP_029760793 | Rdi1 | 97% | 1.00E-52 | 43.94% | Yes |
| Rga1 | XP_029759587 | Rga2 | 33% | 4.00E-28 | 33.33% | Yes |
| Bem3 | XP_029764617 | Bem3 | 21% | 8.00E-87 | 42.69% | Yes |

**Table S1. Polarity proteins identified in *Aureobasidium pullulans*.** Proteins were identified by BLAST using *Saccharomyces cerevisiae* (*S.c.*) proteins as query sequences. RBH: Reciprocal best hit.

| Strain Number | Alt. names | Background | Genotype | Source |
|---|---|---|---|---|
| DLY23540 | EXF-150, ALX001 | | Wildtype EXF-150 | Gostinčar et al., 2014 |
| DLY24148 | ALX002 | DLY23540 | ura3Δ::HYGr | Wirshing et al., 2024 |
| CPY515 | ALX004 | DLY23540 | PAK1-GFP::HYGr | This study |
| CPY516 | ALX005 | DLY23540 | CDC24-GFP::HYGr | This study |
| CPY521 | DLY24601 | DLY23540 | BEM1-GFP::HYGr | This study |
| CPY043 | DLY24148 | DLY24148 | pACT1-3xGFP-BEM1::URA3, ura3Δ::HYGr | This study |
| CPY518 | ALX012 | DLY24148 | CDC42-mCherry(sw)::URA3, ura3Δ::HYGr | This study |
| CPY800 | ALX037 | DLY24148 | RAC1-mCherry(sw)::URA3, ura3Δ::HYGr | This study |
| CPY801 | ALX030 | DLY23540 | pak1Δ::HYGr | This study |
| CPY802 | ALX031 | DLY23540 | cdc24Δ::HYGr | This study |
| CPY803 | ALX032 | DLY23540 | bem1Δ::HYGr | This study |
| CPY804 | ALX033 | CPY043 | pak1Δ::NATr, pACT1-3xGFP-BEM1::URA3, ura3Δ::HYGr | This study |
| CPY805 | ALX036 | CPY043 | PAK1-GFP::HYGr, pACT1-3xGFP-BEM1::URA3, ura3Δ::HYGr | This study |
| CPY806 | ALX027 | CPY515 | cdc42Δ::NATr, Ste20-GFP::HYGr | This study |
| CPY333 | ALX023 | CPY515 | rac1Δ::NATr, Ste20-GFP::HYGr | This study |

**Table S2. Strains used in this study.**

| Description | Parameter | Value |
|---|---|---|
| RD → RT | a | 2 |
| RT → RD | b | 0.5 |
| Diffusion coefficient for RT | Dm | 0.01 |
| Diffusion coefficient for RD | Dc | 10 |

**Table S3. Parameters for the two-component polarity model, related to Figure 2.**

| Description | Parameter | Value |
|---|---|---|
| RD → RT | a | 0.8 |
| RT → RD | b | 0.0875 |
| GEFc → GEFm | c | 0.05 |
| GEFm → GEFc | d | 0.03125 |
| PAKc → PAKm | e | 0.03 |
| PAKm → PAKc | f | 0.04 |
| Diffusion coefficient for RT and GEFm | Dm | 0.01 |
| Diffusion coefficient for RD, GEFc, and PAKc | Dc | 10 |
| Diffusion coefficient for PAKm | Dm2 | 0.1 |

**Table S4. Parameters for the polarity model which includes negative feedback. Related to Figure 5.**

| Description | Parameter | Value |
|---|---|---|
| RD → RT | a | 0.8 |
| RT → RD | b | 0.0875 |
| GEFc → GEFm | c | 0.05 |
| GEFm → GEFc | d | 0.1 |
| Diffusion coefficient for RT and GEFm | Dm | 0.01 |
| Diffusion coefficient for RD, GEFc, and PAKc | Dc | 10 |

**Figure S5. Parameters for the polarity model in which negative feedback was removed. Related to Figure S5.**
